# Cre-mediated deletion of SMARCA5 disrupts pluripotency in mouse embryonic stem cells

**DOI:** 10.1101/2020.04.02.022285

**Authors:** David P. Cook, Barbara C. Vanderhyden

**Author notes:** Correspondence: David Cook, Dr. Barbara Vanderhyden.

## Abstract

In embryonic stem cells (ESCs), the SWI/SNF, CHD, and INO80 families of ATP-dependent chromatin remodellers have been implicated in maintaining pluripotency-associated gene expression. At the time of this study, the importance of ISWI family remodellers had yet to be defined, and we had sought to assess their involvement. During this time, Barisic *et al*. (*Nature*, 2019) elegantly demonstrated that the ISWI homologue SNF2H (*Smarca5*) is important for nucleosomal periodicity, the binding of select transcription factors, and proper differentiation of mouse ESCs. While we do not dispute their findings to any extent, our experiments have led to slightly different conclusions, and we have chosen to use this platform to share our results.

Here, we explore the importance of SNF2H by deriving a conditional knockout mouse ESC line and observing the consequences of SNF2H depletion on the pluripotent state. Cre-mediated deletion of *Snf2h* disrupts hallmark characteristics of pluripotency, resulting in distinct morphological changes; reduced expression of the master transcription factors *Oct4*, *Sox2*, and *Nanog*; and reduced alkaline phosphatase activity. To understand the mechanisms of SNF2H-mediated regulation, we mapped SNF2H-bound nucleosomes genome-wide. SNF2H is broadly distributed across the genome but is preferentially enriched at active regulatory regions and transcription factor binding sites.

## Introduction

Pluripotency is maintained by a coordinated gene expression program that maintains cells in a proliferative, self-renewing state, while preventing the expression of developmental genes. While full length transcripts for approximately half of all protein-coding genes are detectable by RNA sequencing (RNA-Seq) of ESCs^1^, RNA interference (RNAi) screens have repeatedly shown that a subset of several hundred genes are indispensable for the maintenance of a pluripotent state^2–4^. At the core of this network are the transcription factors OCT4, SOX2, NANOG, KLF4, and ESRRB, which form an autoregulatory loop, driving their own expression along with many other actively transcribed genes^5,6^. Chromatin-level regulation also plays a vital role in maintaining ESC identity, involving various histone modifiers, chromatin remodellers, and structural protein complexes. Several of these factors have also been shown to be essential components of the ESC regulatory network, including the Trithorax-group protein WDR5^7^; the histone acetyltransferases P300/CBP^8^; the structural protein CTCF and the Cohesin complex^9,10^; and the nucleosome remodellers BRG1^11^, CHD1^12^, and INO80^13^. Interestingly, when the components of the polycomb repressive complex 2 (PRC2) are deleted in mouse ESCs (mESCs), the expression of pluripotency factors is not affected, however lineage-associated genes become activated and the cells fail to differentiate properly^14^. This is likely due to a failure to maintain repression of developmental genes, as well as failure to repress the mESC gene expression program.

Chromatin remodellers have proven critical for much of mammalian development. Deletion of many individual ATPases, from each of the four families (SWI/SNF, ISWI, CHD, and INO80), precludes the production of a viable embryo, often due to early developmental deficits^15^. Disrupting the expression of many remodeller components in adult cells has highlighted their importance in adult tissues as well. For example, CHD4 is required to maintain the self-renewal of hematopoietic stem cells^16^, and the SWI/SNF ATPase BRG1 is essential for the differentiation into neurons^17^, lymphocytes^18^, adipose^19^, and cardiac tissue^20^. Components of the SWI/SNF, CHD, and INO80 families have each been implicated in maintaining the pluripotent state in mESCs^21–23^, however the ISWI family has yet to be investigated in this context.

The ISWI family of chromatin remodellers in mammals comprises two ATPases, Smarca5 and Smarca1, which are homologues of the *Drosophila melanogaster* ATPase ISWI. Smarca1 expression is limited to reproductive and neural tissues^24^, and has been shown to be important for follicular development within the ovary^25^, and in regulation of progenitor cell populations in the developing brain^26^. Smarca5 is a catalytic subunit of six characterized remodelling complexes (ACF1, CHRAC, RSF, NoRC, WICH, and WCRF) that are broadly involved in transcriptional regulation, DNA replication and repair, and maintaining chromatin structure^27^.

SNF2H also has important roles in mammalian development. It is most studied in the developing cerebellum, where conditional knockout in neural progenitors results in decreased expansion of the progenitor cell population, resulting in neurological deficits and death several weeks after birth^28^. While these cells have reduced proliferation, *Snf2h* deletion in post-mitotic Purkinje cells also produces cognitive alterations, suggesting that observed phenotypes are not solely due to the regulation of proliferation^29^. Its expression is also not limited to neural tissues, but rather, it’s expressed in a variety of tissues^30^. Dynamic *Snf2h* expression during ovarian follicle development also suggests a possible role in regulating proliferating cells in growing follicles^31^. Stopka and Skoultchi^32^ demonstrated that mouse embryos lacking SNF2H are not viable. Using a targeted deletion model that excises a portion of the ATPase domain, they found that homozygous-null embryos are not detectable after 3.5-d.p.c., and when cultured, the inner cell mass of null embryos fails to form a typical colony outgrowth and the cells undergo growth arrest^33^. Heterozygous mice are viable, however. This phenotype is consistent with other null models of remodellers in mice (reviewed by Ho and Crabtree^34^). These data suggest that SNF2H, like other remodellers, may be important in the maintenance of pluripotency and proper development of the early embryo.

## Results

### Derivation of a *Snf2h*^fl/fl^ mESC line

To explore if SNF2H-mediated chromatin remodelling is an essential component of the pluripotency regulatory system, we generated a mESC line from mice with floxed *Snf2h* alleles (*Snf2h*^fl/fl^-mESC), allowing for conditional excision of exon 5 following the addition of Cre recombinase (**Figure 1a**). This excises a portion of the gene encoding the ATPase domain and generates a premature stop codon, preventing expression by nonsense-mediated decay. A knockout system was preferred to knockdown systems because mice harboring only one wild-type *Snf2h* allele are still viable and it is unclear what degree of knockdown would be required to disrupt SNF2H function^32^. Timed matings were performed and blastocysts were recovered at 3.5dpc and cultured on a feeder layer of irradiated MEFs (**Figure 1b**). Cell outgrowths were periodically dissociated, resulting in a cell line with a colony morphology typical of mESCs (**Figure 1c**). PCR-based genotyping confirmed homozygous floxed alleles (**Figure 1d**).

**Figure 1.**
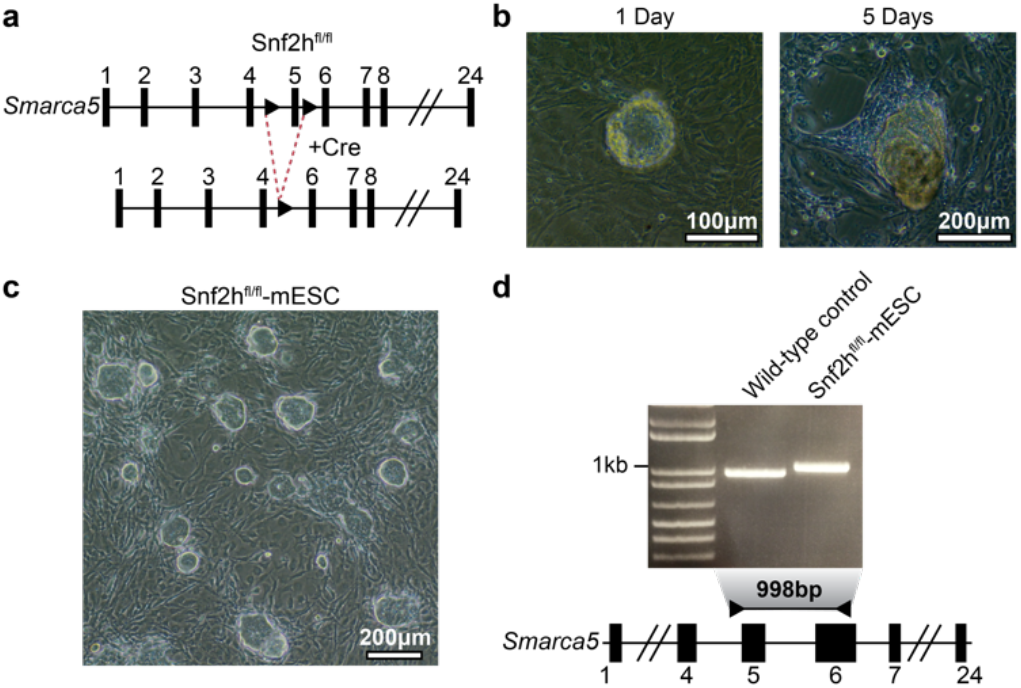
**a)** Schematic representing the Cre-mediated deletion of Snf2h in the *Snf2h*^fl/fl^-mESCs. **b)** Blastocyst collected from a *Snf2h*^fl/fl^ mouse 3.5dpc and cultured on MEF feeders for 1 day (left) and 5 days (right). **c)** Phase contrast image highlighting the morphology of the *Snf2h*^fl/fl^-mESCs. **d)** Gel image confirming Cre-mediated deletion. Gene schematic shows the primer locations used to confirm deletion.

We next validated that the cells displayed characteristics of pluripotency. Expression of the core transcription factors OCT4, SOX2, and NANOG was confirmed by immunofluoresence (**Figure 2a**). When placed in suspension culture, the cells differentiate into embryoid bodies comprising heterogeneous histologies **(Figure 2b**). After approximately 12 days in culture, spontaneously contracting embryoid bodies can be observed. Together, these findings support that the derived *Snf2h*^fl/fl^-mESCs are a bona fide pluripotent mESC line.

**Figure 2.**
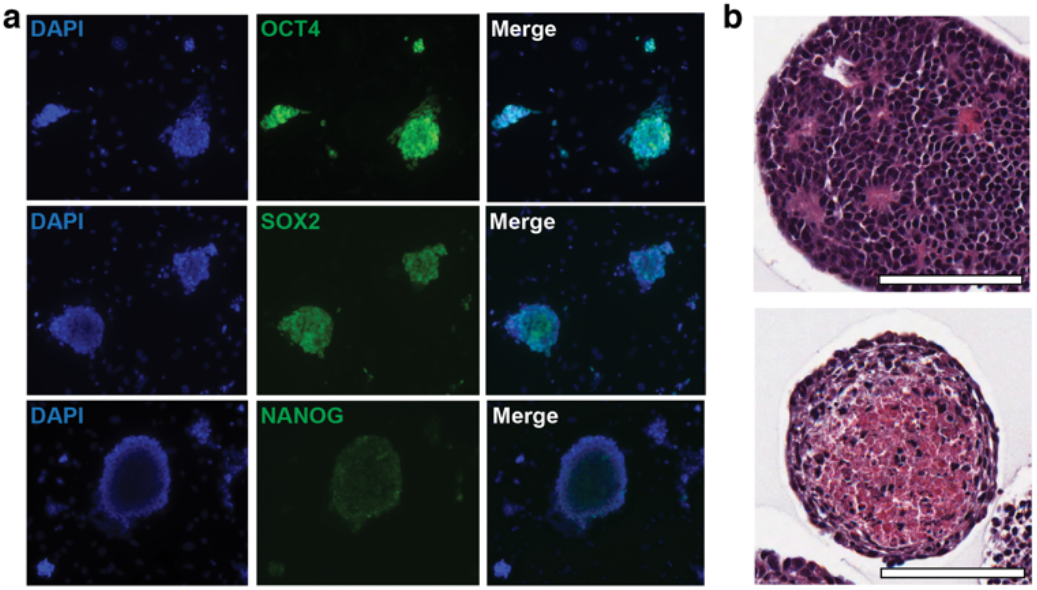
**a)** Immunofluorescence staining showing expression of key pluripotency markers in the *Snf2h*^fl/fl^-mESCs. **b)** H&E stained sections of embryoid bodies that were differentiated for 12 days. Distinct histologies can be observed Scale bars = 100μm.

### Loss of *Snf2h* disrupts pluripotency

To assess the consequences of *Snf2h* deletion on pluripotency, we used a transfection-based approach to introduce Cre recombinase into cells (**Figure 3**). Co-transfection with a GFP vector allowed us to enrich for transfected cells by fluorescence-activated cell sorting (FACS). We found that this was necessary to delay a rebound in the proportion of cells that retained intact *Snf2h* alleles.

**Figure 3.**
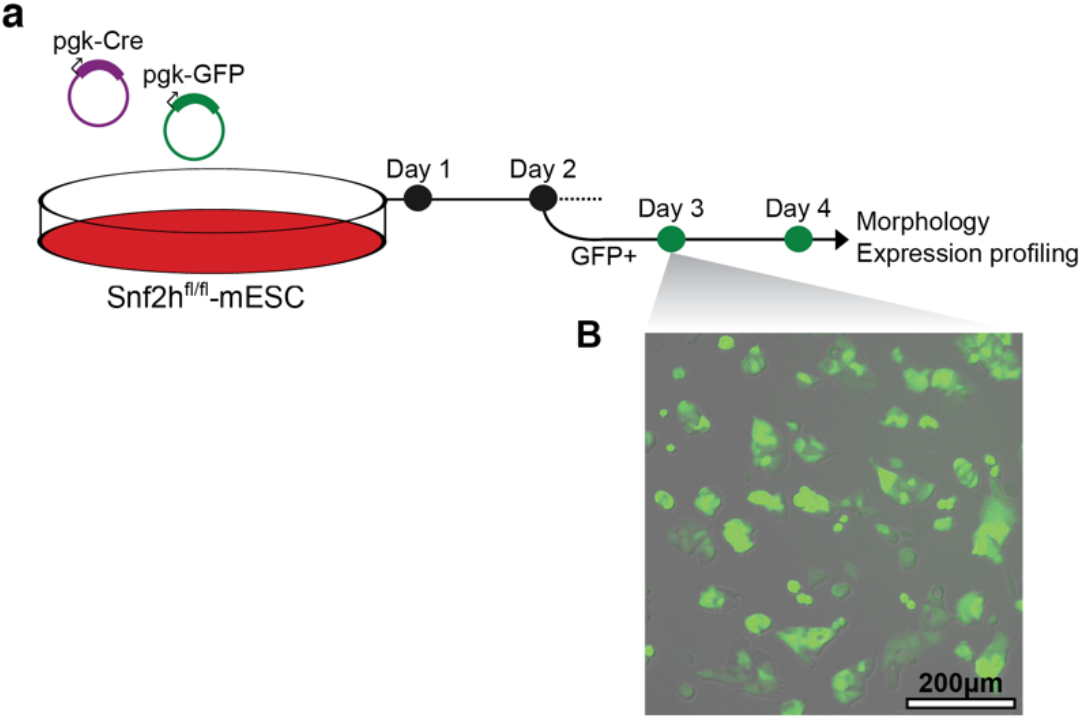
**a)** Transfection strategy used for *Snf2h* deletion. Briefly, cells were cotransfected with Cre and GFP-expressing plasmids. Two days following transfection, GFP+ cells were sorted by FACS to enrich for cells that had been successfully transfected. This reduced heterogeneity from differences in transfection efficiency. Sorted cells were then used for downstream analysis. **b)** Representative population of cells after sorting showing that all cells are GFP+.

Four days following Cre transfection, the cells begin to undergo a marked morphological change, losing their typical colony morphology, flattening out, and spreading across the culture dish (**Figure 4a**). This morphological change is consistent with mESC differentiation, and has been observed following the disruption of other chromatin remodelling enzymes essential to mESCs^21,35,36^. RT-qPCR confirmed an 89% reduction in *Snf2h* expression across the population of cells, confirming that our co-transfection approach yields efficient, albeit not complete deletion throughout the population (**Figure 4b**). Unfortunately, this also showed that *Snf2h* expression levels recover over time, and by 8 days following transfection, are 54% of wild-type levels on average. We also assessed the expression of *Oct4, Sox2*, and *Nanog* and found that all three had reduced expression following the reduction of *Snf2h* expression (**Figure 4b**). The degree of repression was modest; however, this could be explained by both the contamination of *Snf2h-expressing* cells, and the limited time frame assessed.

**Figure 4.**
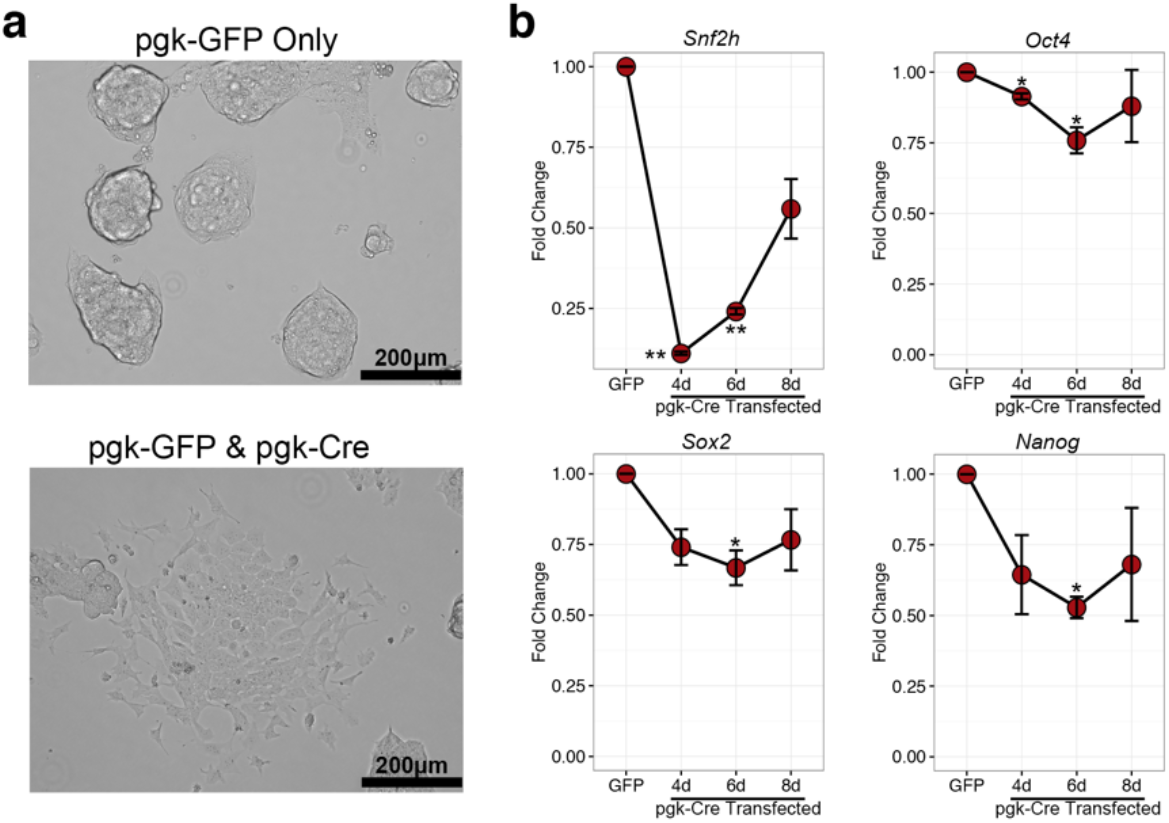
**a)** Brightfield images of GFP-transfected control cells (top) and Cre-transfected cells (bottom). **b)** qPCR gene expression of pluripotency-associated transcription factors and *Snf2h*. Error bars represent SEM. * p-value < 0.05 (One-way ANOVA with Dunnett’s posttest relative to GFP-control cells; n=3).

We also attempted to replicate these findings using a siRNA-based approach in order to confirm that the observed phenotype wasn’t an artifact of our transfection-FACS strategy. We transfected a SMARTpool (Dharmcon) of four unique siRNAs against *Snf2h*, resulting in a 54% knockdown of *Snf2h* levels (**Figure 5a**). Four days following transfection, the cells began to undergo an identical morphological change to Cre-transfected cells (**Figure 5b**). Given the involvement of SNF2H in DNA repair and replication, the observed phenotype may have been the result of cell stress associated with the disruption of these functions. We used an Annexin V assay to assess early apoptosis and cell death at the time of the observed morphological change and found no difference between control and siRNA-transfected cells (**Figure 5c**). This supports that the observed phenotype following *Snf2h* loss is associated with exit from the pluripotent state.

**Figure 5.**
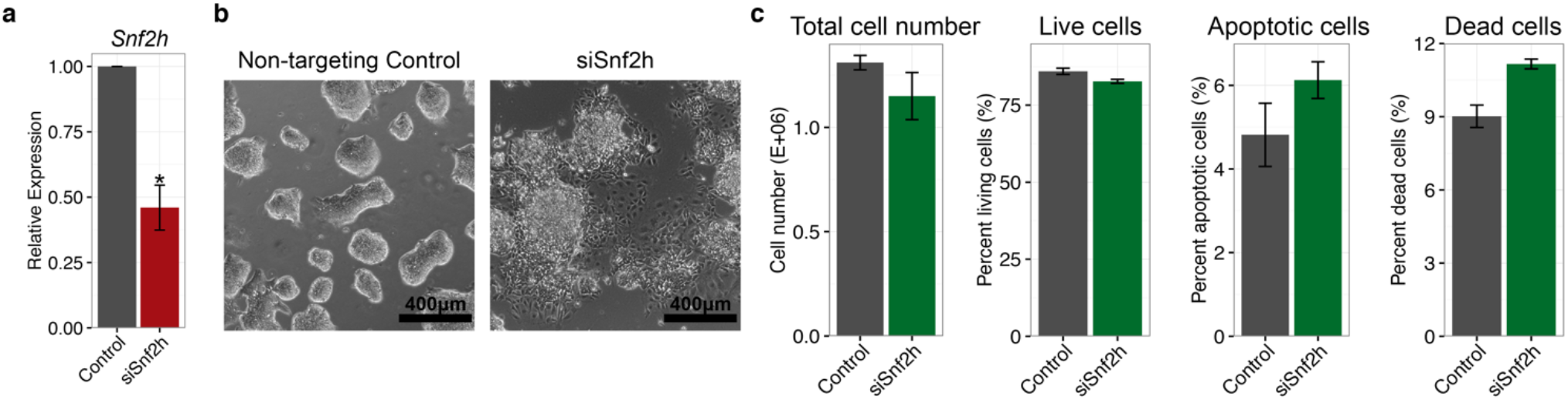
**a)** qPCR expression of *Snf2h* three days following siRNA transfection. Error bars represent SEM. * p-value < 0.05 (Student’s t-test). **b)** Phase contrast images of control (left) and siRNA-transfected (right) cells four days following transfection. **c)** Annexin V and propidium iodide (PI) analysis of the proportions of living (Annexin^low^/PI^low^), apoptotic (Annexin^high^/PI^low^), and dead cells (Annexin^high^/PI^high^). *Snf2h* knockdown does not affect apoptosis/viability at the time of morphological changes (n=3, *p*=0.28, paired two-way ANOVA).

### Snf2h loss activates lineage expression programs

To further support that the *Snf2h* deletion is associated with the loss of pluripotency, we performed RNA sequencing (RNA-Seq) on GFP or Cre-transfected *Snf2h*^fl/fl^-mESCs five days following transfection. The rationale for this time point was to provide enough time following transfection to capture transcriptional consequences of *Snf2h* deletion, while preceding the complete resurgence of wild-type mESCs. *Snf2h* expression was approximately 50% reduced in Cre-transfected samples (**Figure 6a**), suggesting that there was a fair degree of wild-type cells remaining in culture prior to processing. Despite this, however, we found remarkable consistency across biological replicates (**Figure 6b**), allowing us to confidently identify 1445 differentially expressed genes (**Figure 6c**).

**Figure 6.**
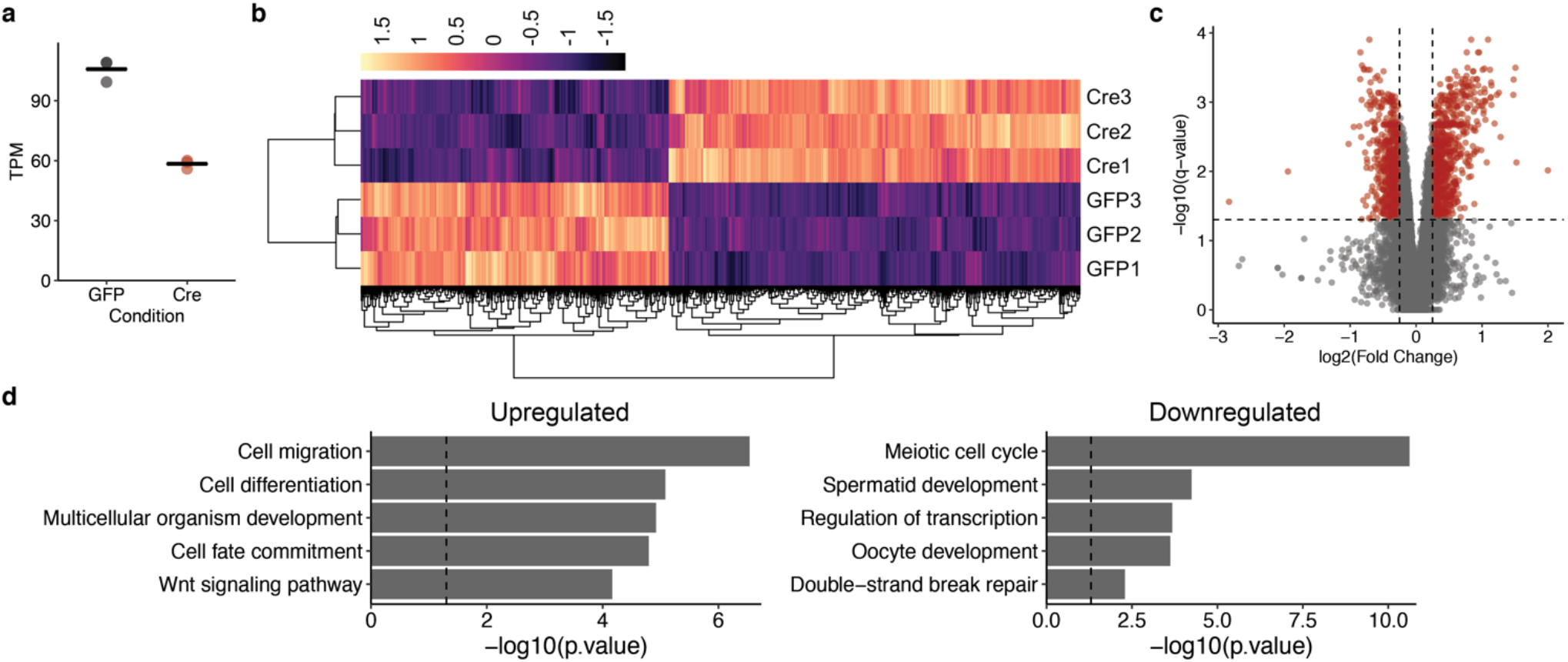
**a)** RNA-Seq expression levels of *Snf2h* across both conditions (Transcripts per million; TPM). Each dot represents a single measurement (n=3) and the straight bar represents the mean. **b)** Heatmap showing the expression (Z-score) of 1445 differentially expressed genes across all samples. **c)** Volcano plot of all genes, showing log2 fold change and *q*-values. Red dots represent genes deemed significantly changed (log2 fold change > 0.25, *q* < 0.05). **d)** Selected GO terms significantly enriched in the upregulated (left) and downregulated (right) genes. The vertical dashed line represents the p-value cutoff of 0.05.

We used gene ontology (GO) term enrichment to assess broad functional associations with the differentially expressed genes. GO terms associated with upregulated genes were generally associated with cell differentiation, proliferation, and various signalling pathways (**Figure 6d**). It was interesting to see such an enrichment of positive regulators of proliferation given the reduced proliferation that enabled resurgence of wild-type mESCs. Downregulated genes interestingly enriched for several terms associated with meiosis and gamete development (**Figure 6d**). Many transcription factors were also downregulated following *Snf2h* deletion, which may be indicative of transitions in the cells’ global gene expression program.

The GO term analysis suggested that *Snf2h* deletion results in the activation of cell differentiation genes, but failed to provide information about which lineage cells differentiate down. We used a previously-curated gene signature for endoderm, ectoderm, mesoderm, and neural lineages^37^ and found that the gene sets for all four lineages were enriched (p < 0.05, Fisher’s exact test) in genes upregulated following *Snf2h* loss. This suggests that deletion of *Snf2h* does not result in a coordinated differentiation program, but rather that the cells randomly differentiate. It’s unclear if each individual cell follows a defined differentiation program, but with heterogeneity across the population of cells, or if *Snf2h* deletion results in uncoordinated transcriptional changes, resulting in the activation of multiple lineages in individual cells.

### Snf2h loss selectively disrupts chromatin accessibility

Chromatin remodelling enzymes are essential for coordinating nucleosome positioning along the genome. We predicted that the loss of pluripotency following *Snf2h* deletion may be the result of disrupted chromatin accessibility. From the same samples that were processed for RNA-Seq, we processed a subset of the cells for ATAC-Seq to assess chromatin accessibility in wild-type and *Snf2h-deleted* cells. While we had originally expected to see dramatic changes in accessibility across the genome, there was surprisingly a high similarity between the accessibility profiles, with a similar number of accessible regions mapped in each condition.

While dramatic changes in accessibility across the genome were not observable, we next narrowed in on the promoter regions of differentially expressed genes to see if *Snf2h* deletion lead to reduced accessibility at downregulated genes. Using the log ratio of ATAC signal from GFP- and Cre-transfected cells to calculate a log enrichment value, we found that accessibility is higher in GFP control cells, suggesting that *Snf2h* deletion results in a loss of chromatin accessibility at these regions (**Figure 7**). The genes that were upregulated following *Snf2h* deletion had a modest increase in chromatin accessibility following Cre transfection (**Figure 7**), but interestingly, these genes also had accessible promoters before Cre transfection. This could be explained by the poised state of the regulatory regions of lineage commitment genes^38^.

**Figure 7.**
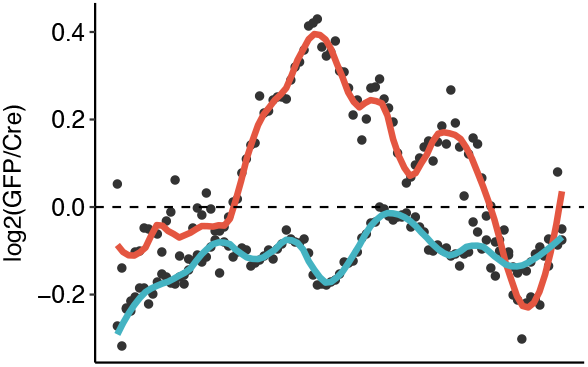
Log2 ratio of GFP-to-Cre ATAC-Seq signal at the promoters of downregulated genes (red) and upregulated genes (blue). The center of the x-axis corresponds to the transcription start site, and the two extremes represent +/− 1kb. Each point represents the averages of specific bins, tiled along the 2kb window. The coloured lines represent LOESS-modelled curves across the 2kb window.

### Snf2h is enriched at active regulatory regions in wild-type mESCs

While this work supports the hypothesis that SNF2H is critical for maintaining pluripotency in mESCs, it doesn’t fully capture its mechanistic involvement in this process. To improve our understanding of this, we performed ChIP-Seq to map the localization of SNF2H across the genome in wild-type *Snf2h*-mESCs. We then assessed the localization of SNF2H at promoters across the genome and found that SNF2H was preferentially enriched at transcriptionally active promoters (**Figure 8a**). Assessing SNF2H ChIP-Seq signal around the binding sites of key mESC transcription factors, we confirmed that it also co-localizes with OCT4, SOX2, and NANOG (**Figure 8b**).

**Figure 8.**
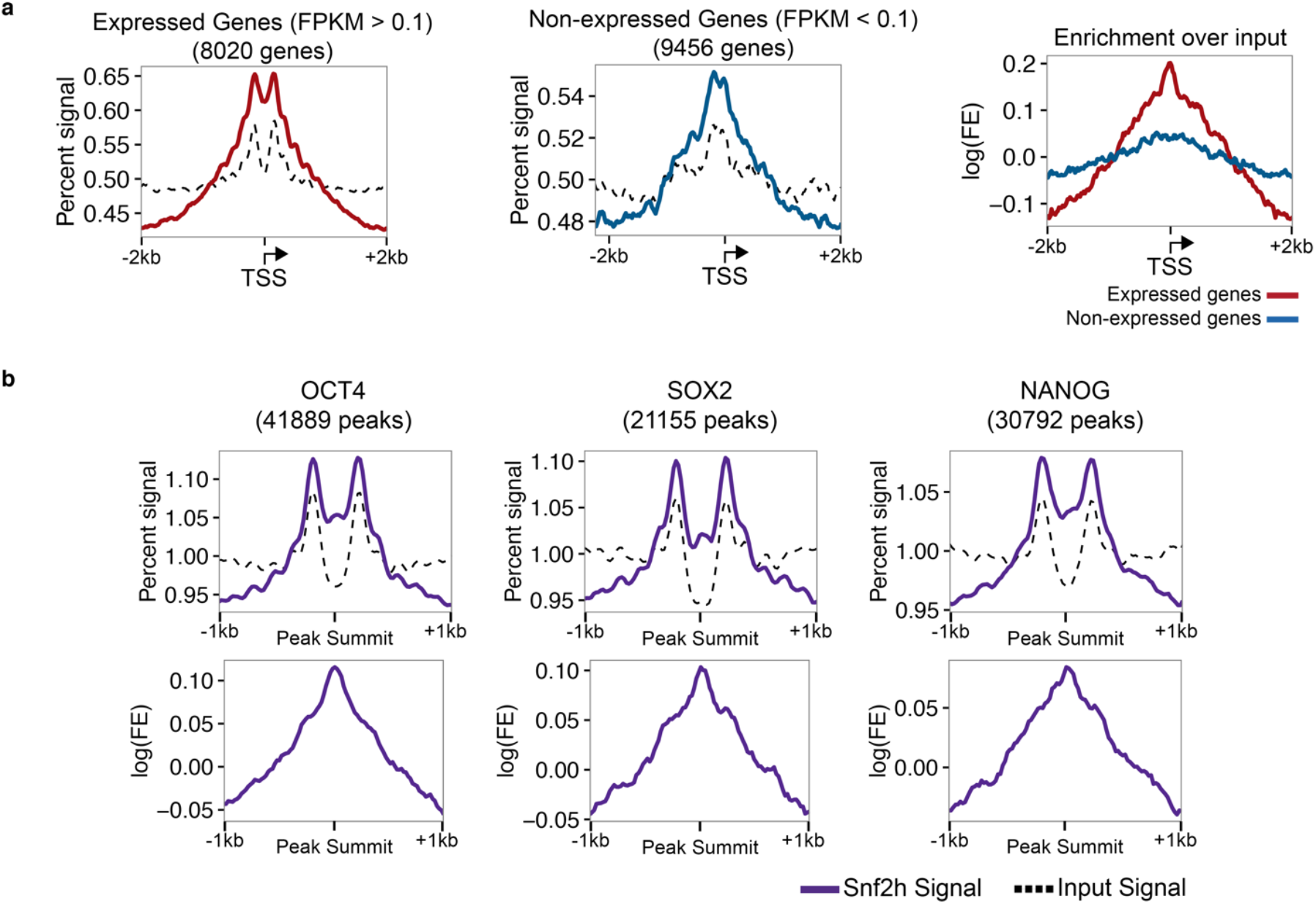
**a)** SNF2H ChIP-Seq signal (coloured line) relative to input controls (dashed line) across the promoters of transcriptionally-active (left) and transcriptionally-repressed (middle) promoters. The right plot compares the log-ratio of the ChIP-Seq signal to input signal at these promoters. **b)** SNF2H ChIP-Seq signal around the binding sites of OCT4 (left), SOX2 (middle), and NANOG (right). The bottom row shows the logratio of ChIP-Seq signal to the input signal.

## Discussion

Here, we show that Cre-mediated deletion of part of SNF2H’s catalytic domain resulted in spontaneous differentiation and decreases in proliferation rates. Shortly following the deletion of SNF2H, we observed upregulation of differentiation-associated genes and a loss of chromatin accessibility at regulatory regions of pluripotency genes, consistent with the more-differentiated morphology. Ultimately, our findings suggest that SNF2H may be critical for the maintenance of pluripotency.

At the time of this study, Barisic *et al*.^39^ demonstrated that generating viable mESCs following CRISPR-mediated deletion of SNF2H was possible and did not affect mESC morphology or expression of pluripotency markers. Rather, its loss lead to altered nucleosomal periodicity, impaired binding of select transcription factors, and failure to form proper embryoid bodies. While some of these findings seem to be in contradiction to our own, we believe they have performed elegant experiments and analyses to come to these conclusions and we do not dispute them. Differences may have arisen from the different experimental approaches, including the strategy to eliminate SNF2H expression. Our study relied on Cre-mediated deletion of exon 5, whereas their line was generated through a CRISPR-induced frameshift in exon 6. Although, we did find that siRNA-mediated knockdown of Snf2h resulted in a similar phenotype. The mESC lines were also from different mouse strains (C57BL/6 in our study, and 129S6/SvEvTac in theirs), which could also contribute to these differences. While the specific findings varied, both studies have shown that SNF2H contributes to regulatory mechanisms in mESCs, adding the ISWI family of chromatin remodellers to the complex regulatory network of pluripotency.

## Methods

### Cell culture and mESC maintenance

mESCs were cultured in standard serum-based culture conditions. Cells were maintained in DMEM with 4.5g/L glucose, L-glutamine, and sodium pyruvate (Corning), supplemented with 15% fetal calf serum, 1X PenStrep (ThermoFisher Scientific), 1X MEM Non-Essential Amino Acids (ThermoFisher Scientific), 1000U/mL ESGRO Leukemia Inhibitory Factor (LIF; EMD Millipore), and 8nL/mL 2-Mercaptoethanol (Sigma). Initial passages of the derived *Snf2h*^fl/fl^-mESCs were maintained on an irradiated feeder layer of DR4 MEFs (ATCC, SCRC1045). After validating characteristics of pluripotency, the line was weaned off feeder cells and cultured on 0.1% gelatinized plates. Culture medium was changed every second day and passaged when confluent (usually every 2-3 days).

### Derivation of a *Snf2h*^fl/fl^-mESC line

The protocol of the use of mice was approved by the University of Ottawa’s Animal Care Committee and was performed in accordance with the guidelines of the Canadian Council on Animal Care. Derivation of the *Snf2h*^fl/fl^-mESC line was performed as described in Czechanski *et al*.^40^. Briefly, timed matings were performed with C57Bl/6 *Snf2h*^fl/fl^ mice^41^ (mice kindly provided by Arthur Skoultchi, Albert Einstein College of Medicine) and 3.5-d.p.c. pregnant females were collected. The uterus was then removed and fertilized embryos were flushed into petri dishes using M2 embryo media (Sigma-Aldrich). Embryos were transferred and washed through several M2 media droplets before being pooled in a petri dish with M2 media. At this stage, most embryos were at the expanded blastocyst stage. Under-developed embryos were transferred into M16 media (Sigma-Aldrich) for further development for up to 24 hours (any embryos that remained under-developed at this point were discarded). No chemical removal of the zona pellucida was performed. Individual hatched blastocysts were then transferred to a well of a 24-well plate containing irradiated DR4 MEF feeders in normal FBS-containing mESC media, described above. Embryos were allowed to attach to the MEF layer for 3 days without disruption. Following attachment, culture medium was changed daily. After 7 days in culture, primary outgrowths were disaggregated using trypsin and vigorous pipetting with a P2 pipette tip. While this tended not to completely break up the primary outgrowth, cells sloughed off in the process did seed the growth of new colonies. After several days (a judgement based on the size of the residual primary outgrowth and the development of secondary colonies), enzymatic disaggregation was repeated. As the number of mESC colonies increased, the cells were eventually transferred to 6-well plates, and finally 10cm plates. The number of disaggregation steps to reach this point was variable, but was approximately 4-6 times. Also, several embryos did not reach this final point, ultimately resulting in two derived lines. One of the two lines was characterized to confirm characteristics of pluripotency, and was used for all experiments.

### PCR-based genotyping for Snf2h deletion

Genomic DNA (gDNA) was isolated from cells using the GeneJET Genomic DNA Purification Kit (ThermoFisher Scientific) and 2μL of purified gDNA was added to 5μL of the 2X REDExtract-N-Amp PCR ReadyMix (Sigma-Aldrich), 0.2μL 10μM forward/reverse primer mix, and 2.8μL ddH_2_O. The reaction was performed with the following parameters: initial denature, 94°C, 3 minutes; denature, 94°C, 30 seconds; anneal, 55°C, 30 seconds; extension, 72°C, 2 minutes; repeat anneal and denature for a total of 34 cycles; one final extension, 72°C, 10 minutes. PCR products were then run on a 1% agarose gel containing RedSafe Nucleic Acid Staining Solution (Intron Biotechnology) and visualized with an EpiChem II Darkroom Transilluminator (UVP Laboratory).

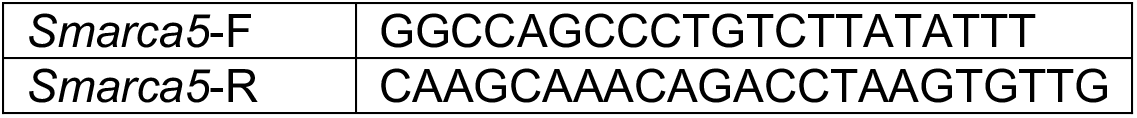
Genotyping primers

### Immunofluorescence of pluripotency factors

Glass coverslips were flame-sterilized, placed in the wells of 6-well plates, and coated with 0.1% gelatin for 30 minutes. The gelatin was then aspirated and *Snf2h*^fl/fl^-mESCs were plated on the gelatinized coverslips. Once the cells had reached confluence, they were washed twice with cold PBS, and then fixed in 4% paraformaldehyde for 30 minutes at room temperature. After fixation, the paraformaldehyde was rinsed off with PBS, and the cover slips were washed three additional times with PBS, bathing the cells for 5 minutes each time. The cells were then permeabilized with 0.2% Triton X-100 in PBS for 10 minutes at room temperature, which was then rinsed off, and the cells were washed for an additional three washes in PBS for 5 minutes each wash. The cells were then blocked in 5% normal goat serum in PBS for 1 hour at room temperature. After blocking, the primary antibodies (OCT4, Ab19857, 1/800; SOX2, Ab97959, 1/1000; NANOG, Ab80892, 1/100) were added and the cells were incubated at 4°C overnight. The following morning, the cells were washed three times in PBS for 5 minutes. The secondary antibody (goat anti-rabbit IgG Alexa Fluor 488, ThermoFisher Scientific, 1/1000) was diluted in 5% normal goat serum in PBS and was added to the cells for 1 hour in the dark, at room temperature. Afterwards, the cells were washed three times for 10 minutes in PBS. Finally, coverslips were removed from the wells and mounted onto slides using ProLong Gold Antifade Mountant with DAPI (ThermoFisher Scientific). Images were taken on a Zeiss Axioskop 2 MOT upright microscope.

### Embryoid body differentiation of *Snf2h*^fl/fl^-mESCs

*Snf2h*^fl/fl^-mESCs (5×10^5^) were diluted into 10mL of mESC media lacking LIF supplementation, and were transferred into a 10cm bacterial petri dish to prevent attachment. Cell aggregates were left alone for 4 days, after which the culture medium was changed every second day. This was performed by transferring the aggregates into a 15mL conical tube, allowing the aggregates to settle at the bottom of the tube, aspirating the cleared media being careful not to aspirate any aggregates, and replacing with 10mL of LIF-free mESC media. This was repeated until 14 days after the initial plating. To prepare the embryoid bodies (EBs) for paraffin embedding and cross-sectioning, the EBs were collected in a 15mL conical tube, the medium was aspirated, and the EBs were washed with PBS. The PBS was then aspirated and the EBs were resuspended in 10% formalin to fix for 30 minutes. Afterwards, the formalin was aspirated, the EBs were washed once in PBS, resuspended in 1mL of 1% agarose, and placed into a square-shaped mold to solidify. After the mix had solidified, the agarose blocks were paraffin-embedded, cut into 5μm sections, and stained with hematoxylin and eosin.

### Co-transfection approach for Cre-mediated deletion of *Snf2h*

*Snf2h*^fl/fl^-mESCs were trypsinized and 3×10^5^ cells were plated per well in a gelatinized 6-well plate. The following day, the culture medium was replaced and the cells were transfected with either pKJ1-GFP alone or a 9:1 ratio of pKJ1-Cre and pKJ1-GFP, respectively (total mass of DNA transfected was the same for both conditions) using Lipofectamine 3000 (ThermoFisher Scientific) according to the manufacturer’s protocol (7.5μL lipofectamine reagent per well). The pKJ1 vectors were a kind gift from Michael McBurney (pKJ1 backbone: Addgene plasmid #11333) and were specifically chosen due to high expression levels in mESCs from the *Phosphoglycerate kinase-1* (*Pgk1*) promoter. We herein refer to these expression vectors as pgk-GFP and pgk-Cre. One day following transfection, GFP expression was visually confirmed and all cells were trypsinized and transferred into a 10cm plate for further expansion. The following day, GFP-expressing cells were purified using FACS. Given that in our experimental group, 9X more pgk-Cre was transfected than pgk-GFP, all GFP-expressing cells should presumably express Cre as well. Sorted cells were then cultured and maintained in 6-well plates for downstream analysis.

### siRNA transfection of *Snf2h*^fl/fl^-mESCs

*Snf2h*^fl/fl^-mESCs were trypsinized and 1×10^5^ cells were plated in a gelatinized 6-well plate. The following day, the culture medium was replaced and the cells were transfected with 75pmol of either siSmarca5 (SMARTpool of four siRNA targeting mouse *Smarca5;* Dharmacon, D-041484-03) or siScrambled (siGenome Non-Targeting siRNA #1; Dharmacon, D-001210-01-05) using Lipofectamine 3000 according to the manufacturer’s protocol for siRNA transfection. Successful knockdown was confirmed by qPCR.

### Quantitative real-time RT-PCR (qPCR)

Cells were trypsinized, centrifuged at 400xg, and washed with PBS before a final centrifugation to isolate a cell pellet. Pellets were frozen at −80°C or immediately processed for RNA extraction on ice. RNA was extracted using the GenElute Mammalian Total RNA Miniprep Kit (Sigma-Aldrich) and RNA quality and quantity were assessed with the NanoDrop ND-1000 spectrophotometer (ThermoFisher Scientific). cDNA was then synthesized from 500ng of RNA using the iScript cDNA Synthesis Kit (Bio-Rad), along with “no reverse transcriptase” and “no RNA” negative controls. Samples were run on an ABI7500 Fast Real-Time PCR System (Applied Biosystems) with the SsoFast EvaGreen Supermix with Low ROX (Bio-Rad). The following parameters were used: initial denature, 95°C, 30 seconds; denature, 95°C, 5 seconds; anneal, 60°C, 30 seconds; repeat denature and anneal steps for a total of 40 cycles.

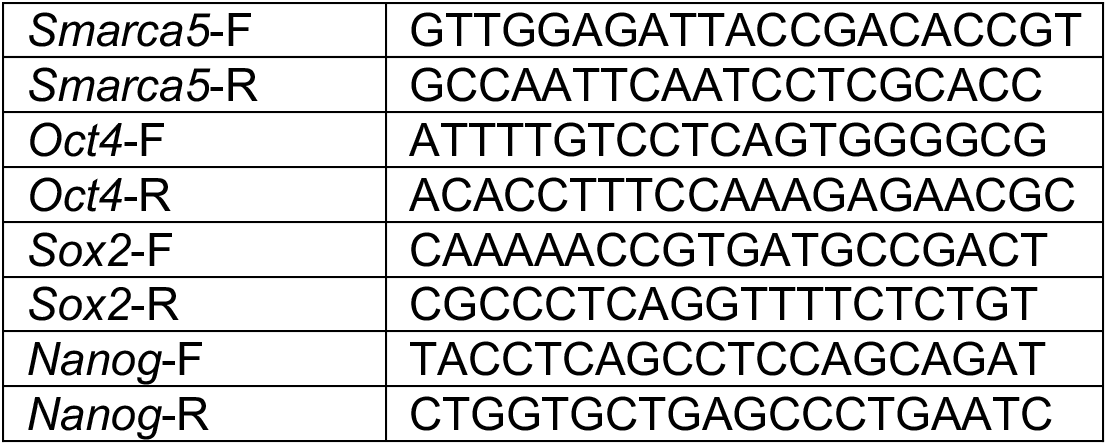
qPCR primer list

### Annexin V apoptosis detection

Annexin V apoptosis and dead cell detection assay was performed with the Alexa Fluor 488 Annexin V/Dead Cell Apoptosis Kit (Invitrogen, cat# V13241) according to the manufacturer’s protocol. Briefly, *Snf2h*^fl/fl^-mESCs were transfected with either siRNA targeting *Snf2h*, or a non-targeting control for mouse cells, as described above, and cultured in normal conditions for four days. Cells were removed from the plate using trypsin, centrifuged at 300xg for 5 minutes, washed with cold PBS, re-centrifuged, and resuspended in 1X annexin-binding buffer at a density of 1×10^6^ cells/mL (100μL per sample). 5μL Alexa Fluor 488 annexin V and 1μL 100μg/mL propidium iodide (PI) solution were added to each 100μL sample. Cells were incubated for 15 minutes in the dark at room temperature. After the incubation, 400μL of 1X annexin-binding buffer was added, mixed gently, and immediately analyzed by flow cytometry. Cells were analyzed by measuring FITC and PI fluorescence. Gates were established to separate three distinct populations of cells: alive cells were FITC^low^ and PI^low^, apoptotic cells were FITC^high^ and PI^low^, and dead cells were FITC^high^ and PI^high^. Three replicates from three independent transfections were performed and identical gates were used across all samples.

### Chromatin immunoprecipitation

Chromatin immunoprecipitation (ChIP) was performed with the SimpleChIP Enzymatic Chromatin IP Kit (Cell Signaling Technology) according to the manufacturer’s protocol. *Snf2h*^fl/fl^-mESCs (20×10^6^) were crosslinked in 1% methanol-free formaldehyde for 10 minutes. Crosslinking was quenched in 1X glycine solution for 5 minutes and cells were washed twice with ice-cold 1X PBS. Cells were scraped from the plate into 1X PBS with 1X protease inhibitor cocktail (PIC) and pelleted by centrifugation at 400xg at 4°C. Nuclei were extracted by resuspending the pellet into 4mL of 1X Buffer A + DTT + PIC for 10 minutes on ice with gentle mixing every 3 minutes. Nuclei were pelleted by centrifugation at 500xg at 4°C, washed in 4mL 1X Buffer B + DTT, re-centrifuged, and resuspended in 400μL 1X Buffer B + DTT. Chromatin was digested using 4μL of micrococcal nuclease (MNase; Cell Signaling Technology, cat #10011) for 25 minutes at 37°C, mixing gently every 5 minutes, producing primarily mono-, and di-, and tri-nucleosomes. The reaction was stopped by adding 40μL of 0.5M EDTA and placing the sample on ice. Nuclei were then pelleted by centrifugation at 15,000xg for 1 minute at 4°C, resuspended in 400μL 1X ChIP Buffer + PIC, and incubated on ice for 10 minutes. In a microcentrifuge tube, nuclei were sonicated for 90 seconds (Misonix Sonicator 3000; 20s ON, 30s OFF) to break the nuclear membrane. Lysates were clarified by centrifugation at 10,000xg for 10 minutes at 4°C.

Clarified lysate (100μL) was transferred to four tubes containing 4μL of 1X ChIP Buffer + PIC and 10μL was removed from one of the tubes and frozen at −20°C for a 2% Input Control. Antibody was added to each tube (10μL anti-Histone H3, Cell Signaling Technology, cat #4620; 4μg Normal Rabbit IgG, Cell Signaling Technology, cat #2729; 4μg anti-SNF2H, Abcam, ab72499). Two SNF2H IP replicates were performed. Samples were mixed with antibodies overnight at 4°C, and 30μL Protein G Magnetic Beads (Cell Signaling Technology, cat #9006) were added for 2 hours at 4°C. Beads were washed three times in low-salt buffer, and once in high-salt buffer. The IP samples were eluted from the beads in 150μL (per IP) 1X ChIP Elution Buffer at 65°C, vortexing every 3 minutes. Crosslinks were then reversed by adding 6μL 5M NaCl and 2μL Proteinase K (Cell Signaling Technology, cat #10012) to each IP sample and incubating at 65°C for 2 hours. DNA samples were column purified (provided by manufacturer), eluted in 50μL DNA Elution Buffer, and quantified using the Quant-iT PicoGreen dsDNA Assay (Invitrogen, cat# P11496).

qPCR was performed by adding 1μL sample with 3μL nuclease-free water, 1 μL 5μM primers, and 5μL SYBR-Green Reaction Mix. The reaction was run with the following parameters: Initial denature, 95°C, 3 minutes; denature, 95°C, 15 seconds; anneal and extension, 60°C, 60 seconds; repeat denature and anneal steps for a total of 40 cycles.

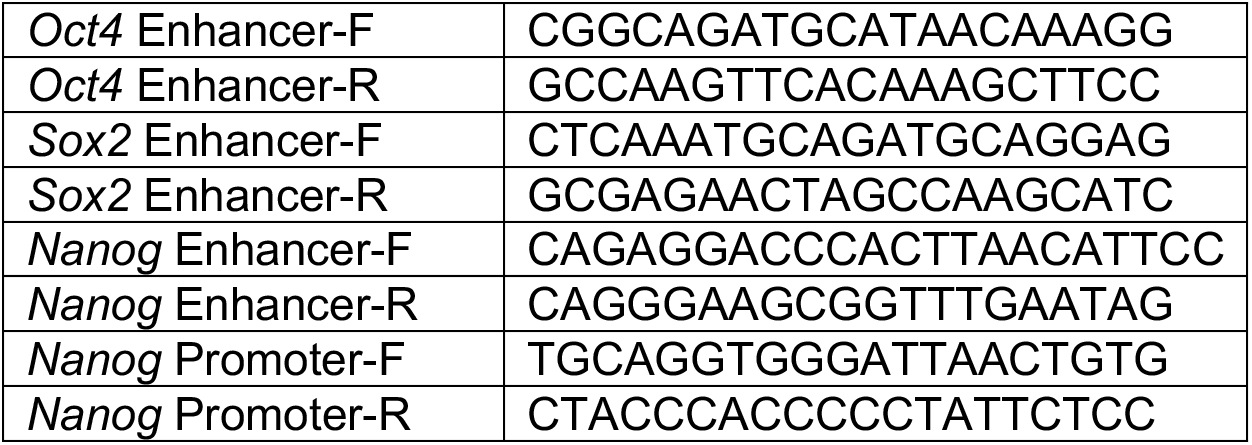
ChIP Primer List

### ChIP-Seq Library preparation and sequencing

Immunoprecipitated and MNase-digested input DNA were submitted to The Centre for Applied Genomics (The Hospital for Sick Children, Toronto, ON) for ChIP-Seq library preparation using the New England Biolabs NEBNext ChIP-Seq Library Prep Reagent Set for Illumina (New England BioLabs) following the manufacturer’s recommended protocol. In brief, DNA was end repaired, followed by dA-tailing to create an overhang A that can ligate to the corresponding overhang T of the double stranded Illumina-compatible adapters, and was PCR enriched with an initial denaturation step at 98°C for 30 seconds, followed by 15 cycles of 98°C for 10 seconds, 65°C for 30 seconds and 72°C for 30 seconds, and a final extension at 72°C for 5 minutes. During adapter ligation, each sample received a different barcoded adapter included in the NEBNext Multiplex Oligos kit (New England BioLabs) to allow for multiplexed sequencing. 1μL of the ChIP-seq libraries was loaded on a Bioanalyzer 2100 DNA High Sensitivity chip (Agilent Technologies) to check for fragment size and absence of primer dimers. Libraries were quantified by qPCR using the Kapa Library Quantification Illumina/ABI Prism Kit protocol (KAPA Biosystems). Libraries were pooled in equimolar quantities and sequenced on an Illumina HiSeq 2500 platform using a Rapid Run Mode flowcell following Illumina’s recommended protocol to generate single end reads of 100 bases in length.

### ChIP-Seq alignment and peak calling

All ChIP-Seq data sets were aligned to the mm9 mouse genome assembly using Bowtie2^42^ with the --sensitive-local parameter. With the exception of the SNF2H ChIP-Seq data sets, peak calling was performed with MACS2^43^ using the callpeak --SPMR -q 0.05 options, along with the -c parameter specifying an appropriate input control when available (See Appendix III for ChIP-Seq data sets and matched controls). These options set a q-value threshold of 0.05 for peak calling and normalize the output signal track by sequencing depth, reporting values as reads per million mapped reads (rpm). Peak calling was omitted for the SNF2H ChIP-Seq data sets because we found it performed poorly at generating high-confidence peak calls. This could be due to its broad distribution across the genome with only modest enrichment at specific loci. Given the high quantification of immunoprecipitated DNA relative to a non-targeting IgG control, we believe this broad signal is true binding rather than technical noise—which is often the interpretation in peak calling—making peak calling inapplicable. Working under the null hypothesis that SNF2H has no regional specificity, binding across the genome with equal probability, evaluating the enrichment of normalized SNF2H signal over MNase-digested input signal at specific target loci proved more useful. For downstream analysis, the alignment files of the two SNF2H ChIP replicates were combined to increase power.

